# Bromelain and Acetylcysteine (BromAc) alone and in combination with Gemcitabine inhibits subcutaneous deposits of pancreatic cancer after intraperitoneal injection

**DOI:** 10.1101/2021.05.05.442745

**Authors:** Ahmad H. Mekkawy, Krishna Pillai, Samina Badar, Javed Akhter, Vahan Képénékian, Kevin Ke, Sarah J. Valle, David L. Morris

## Abstract

**Objective:** Gemcitabine (GEM) is commonly chosen for treating pancreatic cancer. However, its use is limited by toxicity. Earlier in vitro studies with GEM in combination with Bromelain (Brom) and Acetylcysteine (Ac) indicated a substantial reduction in IC50. Here, we investigated the efficacy and safety of Brom and Ac (BromAc) in the pancreatic cancer model in vivo.

**Design:** Both low dose and high dose studies for safety and efficacy of BromAc and GEM were conducted in nude mice. Body weight, wellbeing and tumor volume were monitored. At autopsy, tumor weight, tumor density, percentage of tumor necrosis, expression of Ki67 antigen, and immunohistological evaluation of vital organs were compared between the treatment groups.

**Results:** The low and high doses of BromAc alone and with chemotherapy agents were safe. A very significant reduction in pancreatic tumor volume, weight, and ki67 were seen with BromAc therapy and was equal to treatment with GEM alone and better than treatment with 5-FU. In addition, tumor density was significantly reduced by BromAc.

**Conclusion:** These encouraging results are the first *in vivo* evidence of the efficacy of BromAc in pancreatic cancer and provide some mechanistic leads.

## Introduction

Pancreatic cancer is a lethal malignancy with a very poor prognosis and it is the seventh leading cause of mortality worldwide ^1^, with prediction to be the second by the year 2030 ^2^. In 2018, approximately half a million cases were estimated to be diagnosed with the majority (93%) being fatal ^3^. This dismal outcome has been attributed to late diagnosis owing to non-specific symptoms ^4^. Pancreatic ductal adenocarcinoma (PDAC) is the usual neoplasm in 80-90% of the patients with a median diagnostic age of approximately 70 years ^5^.

Current treatment methods involve radiotherapy, thermo-ablation, surgery, chemotherapy and in some cases only palliation ^6^. Potential for tumor resectability is determined by the absence or presence of distant metastases and locoregional progression – and 90% of patients are not resectable at diagnosis due to their tumor stage ^7^. Over the years a number of chemotherapeutic agents have been used for the treatment of pancreatic cancer such as Gemcitabine (GEM), paclitaxel, 5-FU, Cisplatin, and their combinations have been used with varying success ^8,9^ although more recently FOLFIRINOX (a combination of 5-FU, leucovorin, irinotecan and oxaliplatin) has been advantageous, owing to noticeable survival increase compared to GEM therapy ^10^.

Chemo-resistance accounts for the majority of treatment failures and this has been attributed to the heterogeneity of tumor cells in pancreatic cancer and the tumor extracellular matrix ^11^. Additionally, many molecular alterations have provided chemotherapy resistance to pancreatic cancer ^12^. resistance Mucins have been identified in several cancers including pancreatic cancers ^13^. In PDAC, the expression of several transmembrane mucins and secretory mucins are highly expressed compared to healthy pancreas ^14^. Mucins provide the tumor cells with a barrier defense against drug penetration as well as accelerating survival pathways, chemoresistance, metastasis, and accelerated replication ^15^. Hence, if mucinous barriers can be degraded then there will be an increase in drug penetration resulting in a higher exposure to chemo-agents with better tumor ablation ^16,17^ in addition to abrogating other mucin-enhanced tumor survival pathways. This may lead to a better survival. Pancreatic tumors have very dense tumor matrix that are due to presence of collagen and other proteins including hyaluronic acid leading to vascular damage, decrease tumor perfusion and high Intra Tumoral Fluid Pressure (ITFP) which impairs drug delivery ^18,19^. The collagen present in the intercellular matrix of tumors has both glycosidic and disulfide linkages that are susceptible to the action of certain agents such as Brom and Ac ^20^. Hence, if these barriers to free drug flow can be removed, a better penetration of chemotherapeutic agents into the tumor might be accomplished resulting in a more positive treatment outcome.

Bromelain (Brom) an extract from pineapple (*Ananas Comosus*) fruit or stem contains a number of enzymes such as proteases, carbohydrases, hydroxylases, phosphatases, etc. ^21^ and they have the propensity to hydrolyze the -O- and -N-glycosidic linkages in glycoproteins that are abundant in mucin ^22^. Mucins are polymers of glycoproteins with interlinking disulfide linkages ^23^. In addition, Brom has also shown anti-cancer properties in several studies and currently it is undergoing clinical evaluation for the treatment of mucinous tumors in a rare cancer known as pseudomyxoma peritonei ^24^. Brom is a successful mucolytic in combination with Acetylcysteine (Ac), an antioxidant that can reduce the disulfide bonds found within a mucinous mass ^25^. Ac also has anti-cancer properties in several cancers ^26^. Further, Brom with its proteolytic activities has the ability to disintegrate dense tumor matrix that are made up of collagen, whilst at the same time, it can also disintegrate hyaluronic acid and hence in total allow better passage of drugs into the tumors ^27^.

Remarkably, our previous *in vitro* studies have demonstrated that a suitable combination of Brom with Ac (BromAc) has chemotherapeutic efficacy equivalent to GEM in pancreatic and hepatic carcinoma cells. Noticeably, their synergistic combination with GEM enabled a dramatic reduction of the required dosage of GEM ^28^. BromAc addition to GEM was able to potentiate its efficacy in reduction of colon cancer in vivo ^29^. If the effective dosage of chemotherapeutic agents can be reduced, then, chemotherapy may be given at shorter intervals without increased toxicity. Seven days rest enable resistant or residual tumor cells to replicate and regain potency in the current treatment regime ^30^. Hence, from the encouraging results of our earlier studies ^28^, we proceeded to carry out an *in vivo* evaluation of these agents with safety and efficacy studies in a nude mouse model of pancreatic cancer.

## Materials and Method

### Cell lines

The human pancreatic cancer cell line AsPC-1 was obtained from the American Type Culture Collection and maintained according to the supplier instructions.

### Materials

Bromelain API was manufactured by Mucpharm Pty Ltd (Australia) as a sterile powder. Bromelain was irradiated to ensure sterility. Acetylcysteine was purchased from Link Pharma, Australia (# AUST-R 170803). Gemcitabine hydrochloride was purchased from Sapphire, Australia (Cat # 000-14954, Vend Cat # 1759-25). 5-FU was purchased from Sigma-Aldrich (Cat # F-6627). For treatment, the stock solutions were freshly made and diluted with 0.9% NaCl according to the final treating concentrations required. All other reagents were from Sigma-Aldrich.

### Safety and efficacy study of BromAc in combination with cytotoxics in a nude mice model of pancreatic cancer

Seventy-two, 8-week old female Balb/C nude mice (Animal Resources Center, WA, Australia) were used to examine the efficacy of the combination therapy. After 7 days of acclimatization, 2 × 10^6^ log-phase growing AsPC-1 cells in Matrigel (Cat # E1270, Sigma-Aldrich, Australia) were injected subcutaneous (Day-10). Intraperitoneal treatment was commenced ten days post inoculation to allow the establishment of the disease. Animals were regularly monitored through the study using a standardized method based on the following parameters: body weight, parameters of general wellbeing and indicators of pain and distress classified into four categories (general appearance, natural behavior, provoked behavior and body condition), and tumor volume. Upon completion of the treatment, animals were euthanized, gross appearance of the tumor was examined, tumors were excised, and weighed. The study has been divided into two separate stages: stage 1 (where low doses of GEM and 5-FU have been tested with BromAc) and stage 2 (where high dose of GEM have been tested with BromAc) as follows:

### Safety and efficacy study of BromAc in combination with either gemcitabine or 5-FU in a nude mice model of pancreatic cancer – First stage (Low doses)

Thirsty-six mice (n=6/group) were used in stage 1. Treatments were administered via intraperitoneal injections for 24 days (D1 to D24): BromAc 3 times per week, GEM (2 mg/Kg) and 5-FU (15 mg/Kg) once a week **(Table 1)**. Animals were euthanized on Day 24.

**Table 1:** Efficacy and safety studies of low and high dose treatment regimens. The table shows the type of adjuvant and cytotoxic therapies (GEM or 5-FU) in each treatment group. Treatment was delivered by intraperitoneal route.

| Stage | Group | Therapy Type | Treatment |
| --- | --- | --- | --- |
| Stage 1<br>(Low doses) | T1 | Sham | 0.9% sterile saline solution |
|  | T2 | Combination therapy | Brom (3 mg/kg) + Ac (300 mg/kg) |
|  | T3 | Single-agent therapy | GEM (2 mg/kg) |
|  | T4 | Combination therapy | GEM (2 mg/kg) |
|  |  |  | Brom (3 mg/kg) + Ac (300 mg/kg) |
|  | T5 | Single-agent therapy | 5-FU (15 mg/kg) |
|  | T6 | Combination therapy | 5-FU (15 mg/kg) |
|  |  |  | Brom (3 mg/kg) + Ac (300 mg/kg) |
| Stage 2<br>(High doses) | T1 | Sham | 0.9% sterile saline solution |
|  | T2 | Single-agent therapy | GEM (5 mg/kg) |
|  | T3 | Combination therapy | GEM (5 mg/kg) |
|  |  |  | Brom (3 mg/kg) + Ac (300 mg/kg) |
|  | T4 | Combination therapy | Brom (6 mg/kg) + Ac (500 mg/kg) |
|  | T5 | Combination therapy | GEM (2 mg/kg) |
|  |  |  | Brom (6 mg/kg) + Ac (500 mg/kg) |
|  | T6 | Combination therapy | GEM (5 mg/kg) |
|  |  |  | Brom (6 mg/kg) + Ac (500 mg/kg) |

### Safety and efficacy study of BromAc in combination with gemcitabine in a nude mice model of pancreatic cancer – Second stage (High doses)

Thirty-six mice (n=6/group) were used in stage 2. On day 1, treatment regimen was commenced and continued for another 12 days **(Table 1)**. BromAc was administered every other day (3 times per week; total of 5 doses). GEM (2 or 5 mg/Kg) was administered once/week (total of 2 doses). Animals were euthanized on Day 12.

### Histology and immunohistochemistry

Formalin-fixed, paraffin-embedded sections of tumor as well as various organs were stained using H&E standard techniques. For immunohistochemistry, BOND-III Automated IHC Stainer, Leica has been used. Sections were blocked for non-specific binding, followed by incubation with anti-human Ki67 (Cell Marque; Rabbit Monoclonal Anti-Human; Clone SP6; Cat# 275R-16; Dilution 1/200), incubated with biotinylated anti-rabbit immunoglobulins, treated with streptavidin peroxidase and counter-stained with hematoxylin. The images were captured using a binocular light microscope with a digital camera.

### Statistical analysis

Data were analyzed using GraphPad Prism version 9.0 (GraphPad Software, Inc.). All data were reported as the mean±SD. Qualitative variables were compared using the Student’s *t*-test. Differences were considered statistically significant when *P* < 0.05.

## Results

### In vivo safety and efficacy study – First stage (Low doses)

All animals in the treatment groups survived until 24 days (euthanasia). Their weights increased gradually showing steady growth. No treatment-related toxicities were noted **(Figure 1A)**.

**Figure 1.**
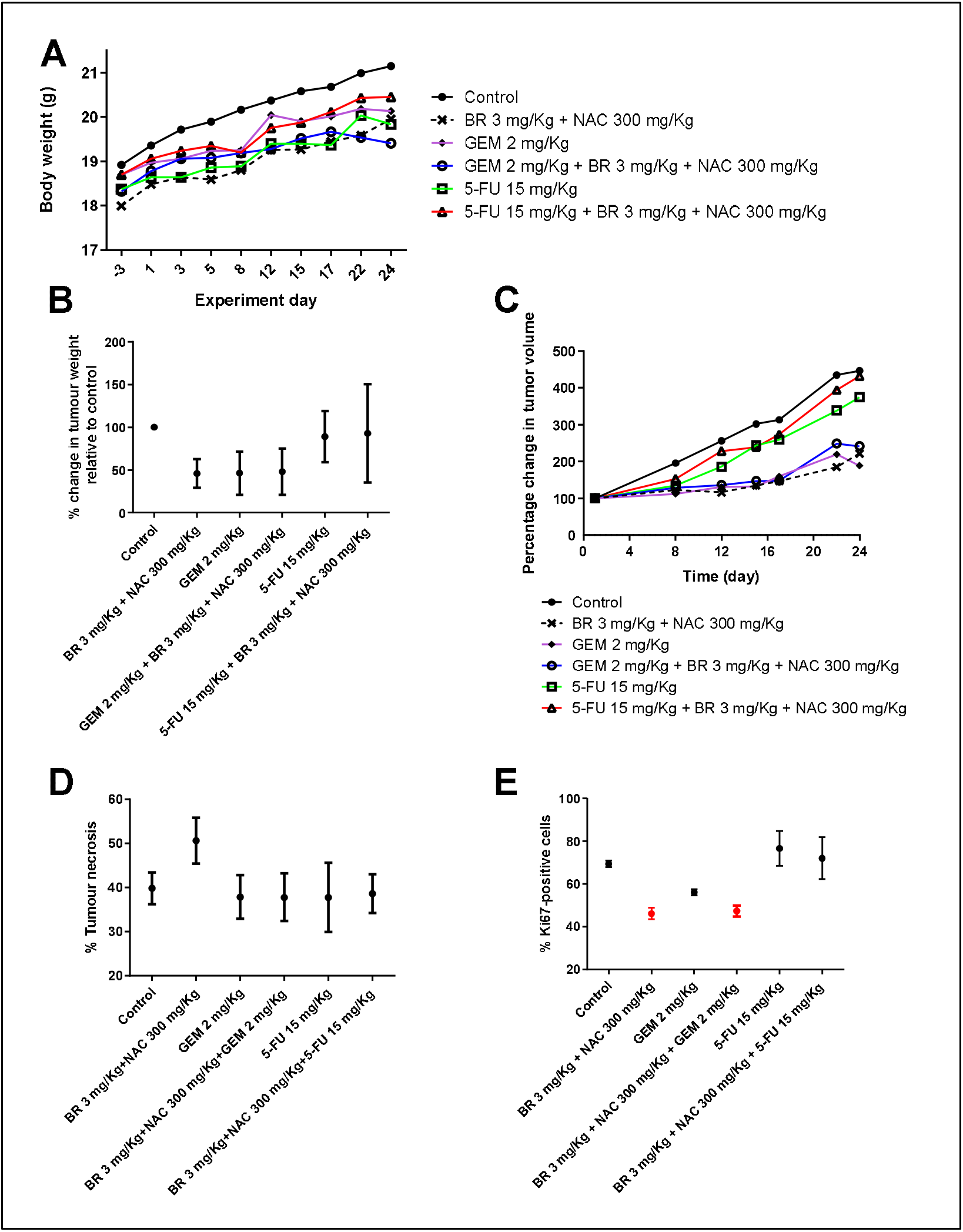
Results of the safety and efficacy in vivo study of BromAc in combination with cytotoxic therapies in AsPC-1 model of pancreatic cancer – First stage (Low doses). **A)** Graph shows mean body weight fluctuations in subcutaneous AsPC1-tumour bearing nude mice treated with combination therapies. **B)** shows percentage change in tumor weight in the treated groups compared to control. **C)** shows percentage change in tumor volume. **D)** Graph showing percentage of tumor necrosis. Necrosis is highest in Brom 3 mg/kg + Ac 300 mg/kg group **E)** Graph showing percentage of Ki-67 positive cells. Analysis of immuno-histological images of tumors samples stained using anti-Ki67 antibody. The lowest expression of Ki67 is observed in groups treated with Brom 3 mg/kg + 300 mg/kg alone or with GEM 2 mg/kg that is indicative of reduced cellular replication. Data presented as mean ± SD.

5-FU alone or with BromAc didn’t inhibit tumor growth based on tumor weight at necropsy **(Figure 1B, Table 2)** whereas BromAc, gemcitabine, or the combination produced a slightly greater than 50% inhibition, indicating a substantial and similar effect on tumor growth.

**Table 2:** Percentage reduction of tumor weight at autopsy in the low dose treatment groups. Percentage reduction of Tumor weight = [Tumor weight (control) – Tumor weight (treatment)/ Tumor weight (control)] × 100. Data are presented as mean ± SD; p-values were obtained through *t*-test; significance level *p* = 0.05.

| **Treatment group** | **Percentage reduction of tumor weight** | **p-value** |
| --- | --- | --- |
| Control |  |  |
| Brom (3.0 mg/kg) + Ac (300 mg/kg) | **54.10±16.72** | **0.0005** |
| GEM (2.0 mg/kg) | **53.55±25.35** | **0.0035** |
| GEM (2.0 mg/kg) + Brom (3.0 mg/kg) + Ac (300 mg/kg) | **51.91±27.33** | **0.0056** |
| 5-FU (15 mg/kg) | 10.93±30.15 | 0.4153 |
| 5-FU (15 mg/kg) + Brom (3.0 mg/kg) + Ac (300 mg/kg) | 7.10±57.53 | 0.7745 |

Tumor volume measurements over 24 days indicated that three treatment groups, Brom (3.0 mg/kg) + Ac (300 mg/kg), GEM (2.0 mg/kg) and GEM (2 mg/kg) + Brom (3.0 mg/kg) + Ac (300 mg/kg) showed hardly any difference from the base line (Day 1) to Day 17, after which the volume increased slightly for the three groups followed by a drop at euthanasia for GEM and BromAc + GEM. Further, the GEM (2 mg/kg) showed a comparatively slightly lower tumor volume **(Figure 1C)**.

When the percentage of tumor necrosis was assessed, it indicated that all the treatment groups except that with Brom 3.0 mg/kg + Ac 300 mg/kg had almost equivalent necrosis (<40%) whilst 50% necrosis was observed in this exceptional group indicating that the combination of Brom and Ac in a weight ratio of 0.01: 1.0 accelerated necrosis **(Figure 1D)**. The mean necrosis was numerically higher in the low BromAc group although results did not reach statistical significance (p = 0.18).

The Ki67 expression in the treatment groups indicated that although three different groups such as Brom 3.0 mg/kg + Ac 300 mg/kg, GEM 2.0 mg/kg and Brom 3.0 mg/kg + Ac 300 mg/kg + GEM 2.0 mg/kg had substantial drop compared to controls, much lower values were observed in the first and the last group indicating the presence of Brom and Ac are crucial for controlling the Ki67 (46%, 56%, 47% respectively compared to control 69%; p < 0.0001) **(Figure 1E)**.

The initial low dose treatment indicated that Brom 3.0 mg/kg + Ac 300 mg/kg, GEM 2.0 mg/kg and Brom 3.0 mg/kg + Ac 300 mg/kg + GEM 2.0 mg/kg had almost equivalent efficacy indicating that any of the dosage forms may be used to derive the present efficacy. This further indicates that at a combination ratio of Brom 3.0 mg/kg + Ac 300 mg/kg, (1:100), no GEM is required for treatment but if GEM is to be added, the dosage can be reduced to an absolute minimum to derive similar maximum therapeutic effect. Hence, this low dose GEM in the presence of Brom and Ac may serve as an effective treatment for pancreatic cancer since more frequent treatment may be instituted (shorter rest interval).

### *In vivo safety and efficacy study* – Second stage (High doses)

Body weight measurement indicated again that all the groups exhibited no negative effect on growth and wellbeing **(Figure 2A)**.

**Figure 2.**
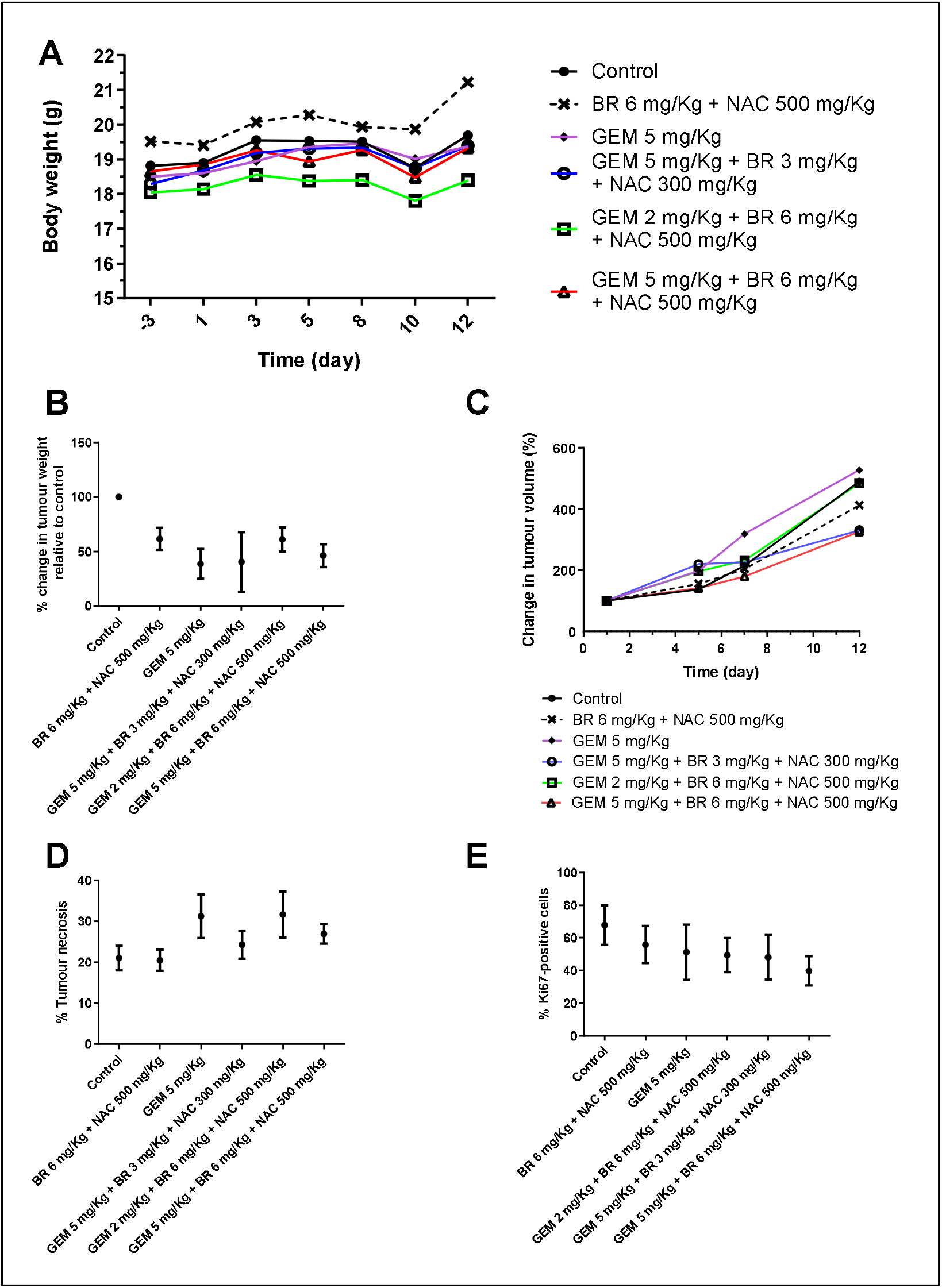
Results of the safety and efficacy in vivo study of BromAc in combination with GEM in AsPC-1 model of pancreatic cancer – Second stage (High doses). **A)** Graph shows mean body weight fluctuations in subcutaneous AsPC-1 -tumour bearing nude mice treated with combination therapies. **B)** shows percentage change in tumor weight in the treated groups compared to control. **C)** shows percentage change in tumor volume. **D)** Graph showing percentage of tumor necrosis. Necrosis is highest in two groups: GEM 5 mg/kg group and Brom 6 mg/kg + Ac 500 mg/kg + GEM 2 mg/kg group **E)** Graph showing percentage of Ki-67 positive cells. Analysis of immuno-histological images of tumors samples stained using anti-Ki67 antibody. The lowest expression of Ki67 is observed in groups treated with Brom 6 mg/kg + 500 mg/kg + GEM 5 mg/kg that is indicative of reduced cellular replication. Data presented as mean ± SD.

When percentage change in tumor weight was assessed at euthanasia, GEM 5.0 mg/kg and GEM 5.0 mg/kg + Brom 3.0 mg/kg +Ac 300 mg/kg showed equivalent and the lowest weight **(Figure 2B)**. This was followed by GEM 5.0 mg/kg + Brom 6.0 mg/kg + Ac 500 mg/kg. **Table 3** shows that GEM at 5.0 mg/kg, and GEM 5 mg/kg + Brom 3.0 mg/kg + Ac 300 mg/kg have an efficacy of 59.57-61.39% in tumor reduction. A difference of 8-10% is observed between the low and the high dose of GEM. The high GEM dose is 2.5 times greater than the low dose.

**Table 3:** Percentage reduction of tumor weight at autopsy in the high dose treatment groups. Percentage reduction of tumor weight = [Tumor weight (control) – Tumor weight (treatment)/ Tumor weight (control)] × 100. Data are presented as mean ± SD; p-values were obtained through *t*-test; significance level *p* = 0.05.

| **Treatment group** | **Percentage reduction of tumor weight** | **p-value** |
| --- | --- | --- |
| Control |  |  |
| Brom (6.0 mg/kg) + Ac (500 mg/kg) | **38.53±10.21** | **0.0048** |
| GEM (5.0 mg/kg) | **61.39±13.69** | **0.0006** |
| GEM (5.0 mg/kg) + Brom (3.0 mg/kg) + Ac (300 mg/kg) | **59.57±27.43** | **0.0225** |
| GEM (2.0 mg/kg) + Brom (6.0 mg/kg) + Ac (500 mg/kg) | **38.94±11.11** | **0.0060** |
| GEM (5.0 mg/kg) + Brom (6.0 mg/kg) + Ac (500 mg/kg) | **53.80±10.52** | **0.0020** |

Tumor volume measurement showed that at Day 7, three groups: (GEM 5.0 mg/kg + Brom 6.0 mg/kg + Ac 500 mg/kg) & (Brom 6.0 mg/kg + Ac 500 mg/kg) showed a lesser tumor volume compared to the rest. However, at Day 12, one of the previously mentioned groups (GEM 5.0 mg/kg + Brom 6.0 mg/kg + Ac 500 mg/kg) in addition to another group (GEM 5.0 mg/kg + Brom 3.0 mg/kg + Ac 300 mg/kg) showed almost equivalent and the lowest tumor volume of the study/ treatments examined **(Figure 2C)**.

Comparison of tumor necrosis to control indicated that Brom 5 mg/kg + Ac 500 mg/kg shared similar outcome, however the other treatment groups showed a higher level of necrosis with groups such as GEM 5.0 mg/kg and GEM 2.0 mg/kg + Brom 6.0 mg/kg + Ac 500 mg/kg showing the highest level of necrosis **(Figure 2D)**.

Comparing the Ki67 expression to control indicated that all treatment groups had a lower level of expression, with GEM 5.0 mg/kg + Brom 6.0 mg/kg + Ac 500 mg/kg expressing the lowest value **(Figure 2E)**.

When the percentage reduction in tumor weights of both the low dose and high dose treatment groups were compared, only two groups (GEM 5.0 mg/kg + Bromelain 3.0 mg/kg + Ac 300 mg/kg & GEM 5.0 mg/kg) showed the highest percentage reduction in tumor weight indicating maximum control over tumor growth.

### Tumor density

Tumor density data are shown in **Table 4 & 5**. It will be seen that the low dose BromAc group (Brom 3mg/kg, Ac 300 mg/kg) produced a 10% reduction in density, 5-FU produced an 8% increase, and in the high dose experiment BromAc produced a 32% decrease in density compared with 34% for GEM alone.

**Table 4:** Density of the tumors from the low dose treatment groups determined at autopsy. Density = tumor mass/ tumor volume (g/cm^3^). Percentage change of tumor density = [Tumor density (control) – Tumor density (treatment)/ Tumor density (control)] x 100.

| **Treatment group** | **Density**<br>**(g/cm<sup>3</sup>)** | **% change**<br>**compared to**<br>**control** |
| --- | --- | --- |
| Control | **0.60** |  |
| Brom (3 mg/kg) + Ac (300 mg/kg) | **0.54** | 10% decrease |
| GEM (2 mg/kg) | **0.59** | 2.0% decrease |
| GEM (2 mg/kg) + Brom (3 mg/kg) +Ac (300 mg/kg) | **0.63** | 5.0% increase |
| 5-FU 15 mg/kg | **0.65** | 8.0% increase |
| **5-FU (15 mg/kg) + Brom (3 mg/kg) + Ac (300 mg/kg)** | 0.60 | No increase |

**Table 5:** Density of the tumors from the high dose treatment groups determined at autopsy. Density = Tumor mass/ Tumor volume (g/cm^3^). Percentage change of tumor density = [Tumor density (control) – Tumor density (treatment)/ Tumor density (control)] x 100.

| Treatment group | Density<br>(g/cm<sup>3</sup>) | % increase or<br>decrease<br>compared to<br>control |
| --- | --- | --- |
| Control | **0.79** |  |
| Brom (6 mg/kg) + Ac (500 mg/kg) | **0.54** | 32% decrease |
| GEM (5 mg/kg) | **0.52** | 34% decrease |
| GEM (5 mg/kg) + Brom (3 mg/kg) + Ac (300 mg/kg) | **0.58** | 27% decrease |
| GEM (2 mg/kg) + Brom (6 mg/kg) + Ac (500 mg/kg) | **0.60** | 24% decrease |
| **GEM (5 mg/kg) + Brom (6 mg/kg) + Ac (500 mg/kg)** | **0.60** | 24% decrease |

### Organ pathology

No histological evidence of abnormality was seen in liver, spleen, kidney, pancreas, and intestine when control and treated groups were compared **(Figure 3)**, indicating complete safety of the different treatment regimens used over the study period.

**Figure 3.**
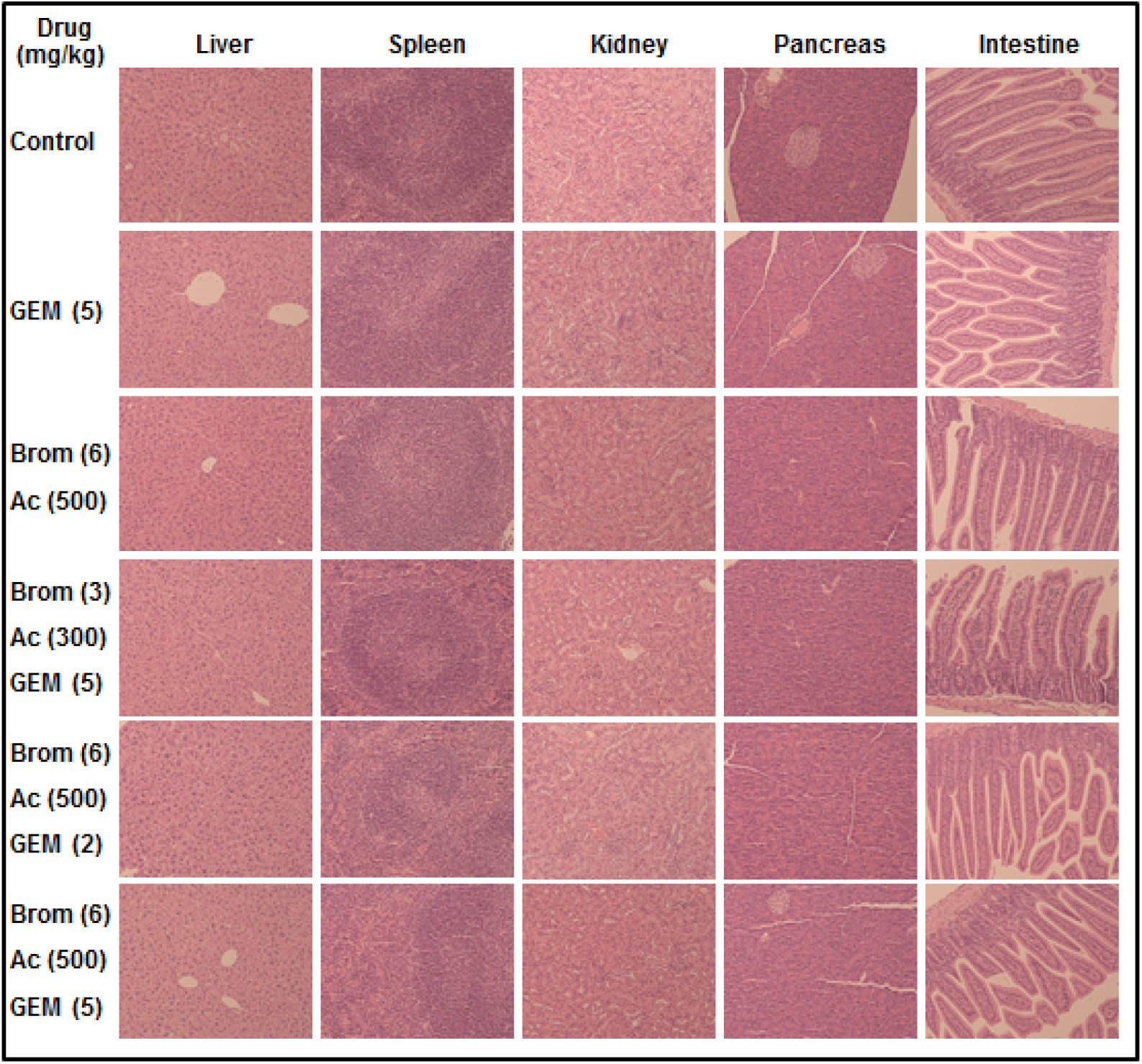
shows the histology of the various organs that may be affected by treatment with GEM, Bromelain and Acetylcysteine and their combinations at various concentrations. Tissues were Hematoxylin and eosin stained (magnification, ×100).

## Discussion

Perhaps the most important finding is that intraperitoneal administration of BromAc can profoundly reduce growth of a xenograft of mucinous pancreatic cancer at a distant subcutaneous site and this is a completely novel observation. Whilst we have previously described in vivo inhibitory effect of BromAc on peritoneal cancers after IP injection ^29^, this paper indicates that a systemic effect of BromAc is possible. The second observation is that BromAc was as effective as GEM in inhibiting tumor growth, and this is the first time that in vivo inhibition of pancreatic cancer growth by BromAc has been reported. We were disappointed that we did not see synergy between BromAc + GEM or 5-FU as seen in our in vitro experiments ^28^. The reason for this disparity deserves further study. The finding that Ki67 was significantly and substantially reduced, and that percentage necrosis was high in the BromAc group supported a strong direct anti-tumor effect. The effect of BromAc on tumor density is also very interesting and may allow better drug penetration in pancreatic cancer.

In the present study the animals tolerated both the low and high dose treatment without any toxicity as revealed by their body weight gain and other parameters of wellbeing, including immuno-histological evaluations of vital organs at termination of the studies. This is a good indication that both low and high dose as stipulated in the treatment regime can be used for clinical application.

Evaluating efficacy through tumor volume and tumor weight regression indicated that in the low dose groups Brom 3.0 mg/kg + Ac 300mg/kg, GEM 2.0 mg/kg and GEM 2 mg/kg + Brom 3.0 mg/kg + Ac 300 mg/kg showed similar efficacy. Further when the percentage change in tumor volume was assessed over the time, the above three groups had similar values indicating that any of the three treatment regimens could be used with equivalent efficacy. The recommended clinical dosage of GEM for pancreatic cancer is 1000 mg/m2 iv ^31^, which is equivalent to 27 mg/Kg ^32^. The low dose of GEM 2 mg/kg (equivalent to a clinical dose of 0.16 mg/kg ^32^) with the addition of BromAc when compared to 27 mg/Kg clinical dose is equivalent to a reduction of more than 99% GEM and it may enable more frequent treatment with foreseeable better tumor ablation and treatment outcome. This minimal GEM dosage regime in the presence of Ac (antioxidant) may reduce the side effects such as neurotoxicity, cardiotoxicity nephrotoxicity etc. ^33,34^. Currently, low dose chemotherapy with gemcitabine is practiced only as palliation ^35^. On the other hand, low dose Bromelain and Ac would allow continuous treatment until tumor is completely ablated. This is a major advantage in using BromAc for treating pancreatic cancer.

Of prime importance is the observation that over treatment period to 17 days, there was almost no tumor growth in the above three treatment groups, indicating the potency of the therapeutic dosage used in controlling the pancreatic cancer growth. Seventeen days translates to almost 2 human years of tumor growth control ^36^ with this treatment strategy. The tumor necrosis was about 50% in the Brom + Ac group (an increase of 10% compared to controls) indicating that these agents besides modulating other oncoproteins may also modulate the vascular epidermal growth factors ^37^ that are responsible for angiogenesis and hence the high level of necrosis that contributes to tumor shrinkage ^38^. Although necrosis is a common feature in most fast-growing tumors, the level in the other groups of treatment was about 40% and similar to controls.

Treatment with 5-FU gave poor outcome as single agent or in combination. Tumor regression was considerably poor and hence further evaluation at high dose was not carried out. The poor outcome with 5-FU may be due to several reasons, being a prodrug, it has to phosphorylated into mono-, di- and tri-phosphorylated fluorouracil compounds ^39^ to be active as a nucleoside. Phosphorylation has been shown to be inhibited by antioxidants such as Ac ^40^.

It is known that tumor matrix are dense owing to several factors such as their composition that are primarily made up of collagen, fibrin fibers hyaluronic acid etc. and with the accumulation of fluid due to leaky blood vessels and poor lymphatic out flow create a very dense environment that restricts the free entry of chemotherapeutic drugs ^41^. In the present treatment the density of the tumor has been evaluated and it was found that treatment groups using Brom 3 mg/kg + Ac 300 mg/kg the density of the tumor fell by 10%, however increasing the dosage to Brom 6 mg/kg + 500 mg/kg Ac, the density was reduced by 32% (a difference of 22%) indicating that Brom and Ac have substantial effect on the tumor matrix and hence the reduction in density. Molecular mechanism that may be at play includes proteolytic action of Brom on collagen, fibrin and other degradable components along with antioxidant action of Ac that has scissoring action on the disulfide bonds linking fibers, proteins etc. ^42^ or the inhibitory effect of Brom on CD44 ^43^ which may alter hyaluronic acid turnover in the tumor stroma. Hence, the present study indicates the safety and efficacy of the treatment regime of BromAc alone or as an adjuvant with GEM as an effective form of treatment for pancreatic cancer.

## DECLARATION OF COMPETING INTEREST

DLM is the co-inventor and assignee of the Licence for this study and director of the spin-off sponsor company, Mucpharm Pty Ltd. AHM, JA, KP and KK are employees of Mucpharm Pty Ltd. SJV is partly employed by Mucpharm for its cancer development and is supported by an Australian Government Research Training Program Scholarship. VK thanks the Foundation Nuovo Soldati for its fellowship and was partly sponsored for stipend by Mucpharm Pty Ltd.

## ETHICS APPROVAL

The animal study was approved by UNSW Animal Care and Ethics Committee (ACEC), Sydney, Australia (approval number: 19/49B).

## FUNDING

This research is partly funded by Mucpharm Pty Ltd, Australia. Grant number: not applicable.

## ACKNOWLEDGEMENTS

We would like to thank Mr John Paul Levi and members of the Pathology Department, St. George Hospital, Kogarah, NSW 2217, Australia for their excellent help in tissue staining.

## Abbreviations

5-FU: 5-Fluorouracil
Ac: Acetylcysteine
Brom: Bromelain
BromAc^®^: Bromelain and Acetylcysteine
CA 19-9: Cancer Antigen 19-9
CI: Combination index
GEM: Gemcitabine
ITFP: Intra Tumoral Fluid Pressure
MUC: Mucin
PDAC: Pancreatic ductal adenocarcinoma.

## Notes

### Competing Interest Statement

The authors have declared no competing interest.

## References

1. Bray F, Ferlay J, Soerjomataram I, et al. Global cancer statistics 2018: GLOBOCAN estimates of incidence and mortality worldwide for 36 cancers in 185 countries. CA Cancer J Clin 2018; 68: 394–424. 2018/09/13.. DOI: 10.3322/caac.21492.

2. Rahib L, Smith BD, Aizenberg R, et al. Projecting cancer incidence and deaths to 2030: the unexpected burden of thyroid, liver, and pancreas cancers in the United States. Cancer Res 2014; 74: 2913–2921. 2014/05/21.. DOI: 10.1158/0008-5472.CAN-14-0155.

3. Siegel RL, Miller KD and Jemal A. Cancer statistics, 2019. CA Cancer J Clin 2019; 69: 7–34. 2019/01/09. DOI: 10.3322/caac.21551.

4. Kamisawa T, Wood LD, Itoi T, et al. Pancreatic cancer. Lancet 2016; 388: 73–85. 2016/02/03. DOI: 10.1016/S0140-6736(16)00141-0.

5. Maire F, Cibot JO, Compagne C, et al. Epidemiology of pancreatic cancer in France: descriptive study from the French national hospital database. European journal of gastroenterology & hepatology 2017; 29: 904–908. 2017/05/05. DOI: 10.1097/MEG.0000000000000901.

6. Matsubayashi H, Kiyozumi Y, Ishiwatari H, et al. Surveillance of Individuals with a Family History of Pancreatic Cancer and Inherited Cancer Syndromes: A Strategy for Detecting Early Pancreatic Cancers. Diagnostics (Basel) 2019; 9: 169. 2019/11/07. DOI: 10.3390/diagnostics9040169.

7. Brunner M, Wu Z, Krautz C, et al. Current Clinical Strategies of Pancreatic Cancer Treatment and Open Molecular Questions. Int J Mol Sci 2019; 20: 4543. 2019/09/22. DOI: 10.3390/ijms20184543.

8. Oettle H, Neuhaus P, Hochhaus A, et al. Adjuvant Chemotherapy With Gemcitabine and Long-term Outcomes Among Patients With Resected Pancreatic Cancer The CONKO-001 Randomized Trial. Jama-Journal of the American Medical Association 2013; 310: 1473–1481. DOI: 10.1001/jama.2013.279201.

9. Neoptolemos JP, Stocken DD, Bassi C, et al. Adjuvant chemotherapy with fluorouracil plus folinic acid vs gemcitabine following pancreatic cancer resection: a randomized controlled trial. JAMA 2010; 304: 1073–1081. 2010/09/09. DOI: 10.1001/jama.2010.1275.

10. Lakatos G, Petranyi A, Szucs A, et al. Efficacy and Safety of FOLFIRINOX in Locally Advanced Pancreatic Cancer. A Single Center Experience. Pathol Oncol Res 2017; 23: 753–759. 2017/01/08. DOI: 10.1007/s12253-016-0176-0.

11. Delitto D, Black BS, Sorenson HL, et al. The inflammatory milieu within the pancreatic cancer microenvironment correlates with clinicopathologic parameters, chemoresistance and survival. BMC cancer 2015; 15: 783. DOI: ARTN 783 10.1186/s12885-015-1820-x.

12. Rajabpour A, Rajaei F and Teimoori-Toolabi L. Molecular alterations contributing to pancreatic cancer chemoresistance. Pancreatology 2017; 17: 310–320.

13. Wang S, You L, Dai M, et al. Mucins in pancreatic cancer: A well-established but promising family for diagnosis, prognosis and therapy. J Cell Mol Med 2020; 24: 10279–10289. 2020/08/04. DOI: 10.1111/jcmm.15684.

14. Krishn SR, Ganguly K, Kaur S, et al. Ramifications of secreted mucin MUC5AC in malignant journey: a holistic view. Carcinogenesis 2018; 39: 633–651. 2018/02/08. DOI: 10.1093/carcin/bgy019.

15. Weniger M, Honselmann KC and Liss AS. The Extracellular Matrix and Pancreatic Cancer: A Complex Relationship. Cancers (Basel) 2018; 10 2018/09/12. DOI: 10.3390/cancers10090316.

16. Choi IK, Strauss R, Richter M, et al. Strategies to increase drug penetration in solid tumors. Front Oncol 2013; 3: 193. 2013/07/31. DOI: 10.3389/fonc.2013.00193.

17. Dauer P, Nomura A, Saluja A, et al. Microenvironment in determining chemo-resistance in pancreatic cancer: Neighborhood matters. Pancreatology 2017; 17: 7–12.

18. Neesse A, Bauer CA, Öhlund D, et al. Stromal biology and therapy in pancreatic cancer: ready for clinical translation? Gut 2019; 68: 159–171.

19. Jacobetz MA, Chan DS, Neesse A, et al. Hyaluronan impairs vascular function and drug delivery in a mouse model of pancreatic cancer. Gut 2013; 62: 112–120.

20. Shoulders MD and Raines RT. Collagen structure and stability. Annu Rev Biochem 2009; 78: 929–958. 2009/04/07. DOI: 10.1146/annurev.biochem.77.032207.120833.

21. Harrach T, Eckert K, Schulze-Forster K, et al. Isolation and partial characterization of basic proteinases from stem bromelain. J Protein Chem 1995; 14: 41–52. 1995/01/01. DOI: 10.1007/BF01902843.

22. Bansil R and Turner BS. Mucin structure, aggregation, physiological functions and biomedical applications. Current Opinion in Colloid & Interface Science 2006; 11: 164–170. DOI: 10.1016/j.cocis.2005.11.001.

23. Meldrum OW, Yakubov GE, Bonilla MR, et al. Mucin gel assembly is controlled by a collective action of non-mucin proteins, disulfide bridges, Ca(2+)-mediated links, and hydrogen bonding. Sci Rep 2018; 8: 5802. 2018/04/13. DOI: 10.1038/s41598-018-24223-3.

24. Valle SJ, Akhter J, Mekkawy AH, et al. A novel treatment of bromelain and acetylcysteine (BromAc) in patients with peritoneal mucinous tumours: A phase I first in man study. Eur J Surg Oncol 2021; 47: 115–122. 2019/11/05. DOI: 10.1016/j.ejso.2019.10.033.

25. Pillai K, Akhter J, Chua TC, et al. A formulation for in situ lysis of mucin secreted in pseudomyxoma peritonei. Int J Cancer 2014; 134: 478–486. 2013/07/12. DOI: 10.1002/ijc.28380.

26. Aluigi MG, De Flora S, D’Agostini F, et al. Antiapoptotic and antigenotoxic effects of N-acetylcysteine in human cells of endothelial origin. Anticancer Res 2000; 20: 3183–3187. 2000/11/04.

27. Parodi A, Haddix SG, Taghipour N, et al. Bromelain surface modification increases the diffusion of silica nanoparticles in the tumor extracellular matrix. ACS Nano 2014; 8: 9874–9883. 2014/08/15. DOI: 10.1021/nn502807n.

28. Pillai K, Mekkawy AH, Akhter J, et al. Enhancing the potency of chemotherapeutic agents by combination with bromelain and N-acetylcysteine - an in vitro study with pancreatic and hepatic cancer cells. Am J Transl Res 2020; 12: 7404–7419. 2020/12/15.

29. Mekkawy A, Pillai K, Badar S, et al. Addition of bromelain and acetylcysteine to gemcitabine potentiates tumor inhibition in vivo in human colon cancer cell line LS174T. Am J Cancer Res 2021; 11.

30. Hegde GV, de la Cruz C, Eastham-Anderson J, et al. Residual tumor cells that drive disease relapse after chemotherapy do not have enhanced tumor initiating capacity. PloS one 2012; 7: e45647.

31. Gemzar, Highlights of Prescribing Information, The food and Drug Administration (FDA), Revised: 05/2019, https://www.accessdata.fda.gov/drugsatfda_docs/label/2005/020509s033lbl.pdf, (Accessed on 03 May 2021).

32. Nair AB and Jacob S. A simple practice guide for dose conversion between animals and human. Journal of basic and clinical pharmacy 2016; 7: 27.

33. Hilmi M, Ederhy S, Waintraub X, et al. Cardiotoxicity Associated with Gemcitabine: Literature Review and a Pharmacovigilance Study. Pharmaceuticals (Basel) 2020; 13: 325. 2020/10/25. DOI: 10.3390/ph13100325.

34. Hryciuk B, Szymanowski B, Romanowska A, et al. Severe acute toxicity following gemcitabine administration: A report of four cases with cytidine deaminase polymorphisms evaluation. Oncol Lett 2018; 15: 1912–1916. 2018/02/13. DOI: 10.3892/ol.2017.7473.

35. Zwitter M, Kovac V, Smrdel U, et al. Gemcitabine in brief versus prolonged low-dose infusion, both combined with cisplatin, for advanced non-small cell lung cancer: a randomized phase II clinical trial. J Thorac Oncol 2009; 4: 1148–1155. 2009/06/24. DOI: 10.1097/JTO.0b013e3181ae280f.

36. Dutta S and Sengupta P. Men and mice: Relating their ages. Life Sci 2016; 152: 244–248. 2015/11/26. DOI: 10.1016/j.lfs.2015.10.025.

37. Redondo P, Bandres E, Solano T, et al. Vascular endothelial growth factor (VEGF) and melanoma. N-acetylcysteine downregulates VEGF production in vitro. Cytokine 2000; 12: 374–378. 2000/05/11. DOI: 10.1006/cyto.1999.0566.

38. Vayrynen SA, Vayrynen JP, Klintrup K, et al. Clinical impact and network of determinants of tumour necrosis in colorectal cancer. Br J Cancer 2016; 114: 1334–1342. 2016/05/20. DOI: 10.1038/bjc.2016.128.

39. Miura K, Kinouchi M, Ishida K, et al. 5-fu metabolism in cancer and orally-administrable 5-fu drugs. Cancers (Basel) 2010; 2: 1717–1730. 2010/01/01. DOI: 10.3390/cancers2031717.

40. Fu Y, Yang G, Zhu F, et al. Antioxidants decrease the apoptotic effect of 5-Fu in colon cancer by regulating Src-dependent caspase-7 phosphorylation. Cell Death Dis 2014; 5: e983. 2014/01/11. DOI: 10.1038/cddis.2013.509.

41. Senthebane DA, Jonker T, Rowe A, et al. The Role of Tumor Microenvironment in Chemoresistance: 3D Extracellular Matrices as Accomplices. Int J Mol Sci 2018; 19 2018/09/23. DOI: 10.3390/ijms19102861.

42. Aldini G, Altomare A, Baron G, et al. N-Acetylcysteine as an antioxidant and disulphide breaking agent: the reasons why. Free Radic Res 2018; 52: 751–762. 2018/05/11. DOI: 10.1080/10715762.2018.1468564.

43. Harrach T, Gebauer F, Eckert K, et al. Bromelain proteinases modulate the cd44 expression on human molt-4/8 leukemia and sk-mel-28 melanoma-cells in-vitro. Int J Oncol 1994; 5: 485–488. 1994/09/01.

